# Reduced dimension stimulus decoding and column-based modeling reveal architectural differences of primary somatosensory finger maps between younger and older adults

**DOI:** 10.1101/2023.01.30.526109

**Authors:** Avinash Kalyani, Oliver Contier, Lisa Klemm, Elena Azañon, Stefanie Schreiber, Oliver Speck, Christoph Reichert, Esther Kuehn

## Abstract

The primary somatosensory cortex (SI) contains fine-grained tactile representations of the body, arranged in an orderly fashion. Using ultra-high resolution fMRI data to describe such detailed individual topographic maps or to detect group differences is challenging, because group alignment often does not preserve the high spatial detail of the data. Here, we use shared response modeling (SRM), a technique that allows group analyses by mapping individual stimulus-driven responses to a lower dimensional shared feature space, to detect age-related differences in sensory representations between younger and older adults using 7T-fMRI data. Using this method, we show that finger representations are more precise in Brodmann-Area (BA) 3b and BA1 compared to BA2 and motor areas, and that this hierarchical processing is preserved across age groups. By combining SRM with column-based decoding (C-SRM), we further show that the number of columns that optimally describes finger maps in SI is higher in younger compared to older adults in BA1, indicating a greater columnar size in older adults’ SI. Taken together, we conclude that SRM is suitable for finding fine-grained group differences in SI fMRI data at ultra-high-resolution, and we provide first evidence that the columnar architecture of a functional area changes with increasing age.

## 1 Introduction

Data aggregation is vital to increase the statistical power of analyses across individuals with varying functional topographies (Dervin, 1990; Mazziotta et al., 2001). For example, whereas differences in the functional topography of the hand area in sensory cortex between younger and older adults are almost impossible to detect at the single-subject level, group analyses allow detecting reliable and consistent differences across age groups (Liu et al., 2021). The study of multiple subjects is therefore crucial to comment on the significance of functional response differences generated in response to a common stimulus. A common problem, however, is inter-subject data alignment, particularly of ultra-high-field magnetic resonance imaging (UHF-MRI) data at a field strength of 7 Tesla (T) or above, where detailed features are encoded in a much higher spatial resolution, which sometimes does not align with the features used for anatomical alignment. Precise alignment is needed, however, to, for example, compare different age groups, compare patients against controls, or investigate changes across different time points in the course of learning or aging.

Functional and anatomical inter-subject alignment are two methods to create a common analysis space (Sabuncu et al., 2010), where anatomical alignment uses anatomical landmarks, such as gyri or sulci, to achieve this goal (Brett et al., 2002; Fischl et al., 1999). Functionally defined regions are not necessarily consistent with these anatomical landmarks. For example, the location of the area related to visual motion perception (area MT) can show a variation of around 2 cm after anatomical normalization (Sabuncu et al., 2010). There are different reasons why anatomical normalization often fails to capture the fine-grained functional differences between individuals’ brains, in particular, when UHF-MRI data is used: On the one hand, smoothing across individuals is required as a step of normalization, and on the other hand, anatomical landmarks may not coincide with the structure of functional circuits (Guntupalli et al., 2016), making it inefficient for UHF-MRI inter-subject analysis.

Functional magnetic resonance imaging (fMRI) data is complex by nature. If every voxel represents a dimension of variation, the activity patterns over these voxels can be described as data points in a high dimensional space. The large number of voxels and the smaller number of volumes (TRs or time points) makes the high dimensional space sparse by nature, due to which statistical analyses are difficult and underpowered. Shared response modeling (SRM) (Chen et al., 2015) is a possible solution for the problems outlined above because SRM projects the fMRI time series of each participant to a low-dimensional space, which captures the temporal variance shared across participants when exposed to the same stimulus or task sequence (for example: watching a movie (Häusler and Hanke, 2021)). The experimental manipulation or stimulus induces a series of cognitive or sensory states, like visual, auditory, or semantic, and shared variance is used to highlight the common variance related to these specific states between participants. SRM, therefore, improves the statistical power by aligning fine-grained spatial patterns in the specific region of interest (Cohen et al., 2017). Even though this technique has great potential to solve common problems in UHF-MRI analyses, it has rarely been used for this purpose so far. Another similar technique that improves the alignment of fMRI time series across participants is hyperalignment (Guntupalli et al., 2016, (also see Feilong et al., 2021, 2018)). It uses a Procrustean transformation to align the activity pattern across individuals into a higher dimensional common model space. For example, Kilmarx et al. (2021) used hyperalignment to align functional time series across individuals corresponding to finger presses, and then performed leave-one-subject-out classification. They found a significant improvement in classification accuracy (p < 0.001) of individual finger presses when group data was aligned based on function (88%) rather than anatomy (46%).

In a similar study by Al-Wasity et al. (2020), the authors successfully predicted imagined arm movements in the motor cortex using techniques of hyperalignment. While both studies involved sensorimotor tasks specific to digits and hands, in the present work, we focus on the fine-grained activity patterns evoked by passive touch in younger and older adults. To reach this goal, we use high-resolution fMRI with an isotropic resolution of 1 mm, acquired at a 7 Tesla MRI scanner.

In this study, we focus on the primary somatosensory cortex (SI) as a model system, whereas our outlined analysis approaches can also be applied to other brain areas or functional units. SI encodes tactile input to the skin in multiple subregions, in particular Brodmann Area (BA)1, BA2, and BA3b. Prior studies have shown that the topographic architecture in somatosensory representations differs in these subareas (Cassady et al., 2020). In addition, higher representational overlap is expected in older adults compared to younger adults (Cabeza, 2002; Liu et al., 2021). However, in most studies, either 3T-fMRI data was used, or when 7T-fMRI data was used, functional features such as overlap or representational similarity were first extracted at the individual subject level and then averaged across the group, preventing voxel-wise analyses across the group. A further open aspect about the functional organization of SI in younger and older adults is its columnar architecture. Due to recent advances in fMRI techniques, we can now measure highly sensitive and spatially accurate signals, making it possible to study fundamental computational units (Yacoub et al., 2008) such as cortical columns. Two good examples of such a functional columnar organization in the brain that have already been studied using fMRI are the orientation columns in the primary visual cortex (V1) (Erwin et al., 1995), and fine-grained movement-dependent finger maps in the primary motor cortex (MI) ((Huber et al., 2017, 2020a), see (Kuehn and Pleger, 2020) for review). For SI, it is unclear whether or not such columnar analyses can be used to describe the system, which columnar size would best represent SI finger maps, and whether columnar sizes would differ between younger and older adults. The question of whether or not columnar sizes differ with age is an aspect that has generally not received much attention in the literature so far. Please note that the term ‘columnar’ here refers to the mesoscopic structures aligned perpendicular (radial) to the cortical depths, in the context of fMRI data (Huber et al., 2020a).

Here, we use SRM to obtain information about the functional architecture of sensory finger representations in the SI of younger and older adults. More specifically, we investigate decoding sensitivity for finger discrimination across different BAs (BA3b, BA1, BA2) in SI as well as across the motor system (BA4a, BA4p, BA6), and compare them between younger and older adults. Then, we introduce a column-driven SRM approach where the number of columns is varied as a hyperparameter to differentiate the vibrotactile stimulation of fingers. In addition, we test whether this hyperparameter varies across age groups. Using the same principle of columnar mapping as introduced above, we study here for the first time these computational units in SI to determine the optimal columnar scale to describe finger representations in columnar units in younger and older adults. We argue that this method can in the future be also applied to other brain areas to determine the optimal columnar size within which functional maps can be described (for example in parietal cortex, insula, and hippocampus). The columns we describe here may represent the smallest unit of functional processing, i.e., population receptive fields. We further argue that the set of techniques we offer here provides key information about the architecture of SI, and is particularly useful for analyzing UHF-MRI data where the preservation of the individual information and spatial resolution is a key challenge.

## 2 Methods

### 2.1 Participants

We tested m = 19 younger adults (mean age 25 ± 0.49, ranging from 21 to 29 years, number of females = 9) and m = 19 older adults (mean age 72.2 ± 0.81, ranging from 65 to 78 years, number of females = 9). These participants were recruited from the database of the DZNE Magdeburg. They were all healthy and were checked for 7T-MRI exclusion criteria, such as metallic implants and other foreign bodies, active implants (e.g. pacemaker, neurostimulator, cochlear implant, defibrillator, and pump system), permanent makeup, tinnitus, or other hearing impairments. The functional 7T-MRI scan was performed in a single session alongside a whole-brain MP2RAGE sequence, and one additional 3T-MRI session took place on a separate day (see below). All participants were compensated for their time and attendance, and a written informed consent form was signed by the participants before each scan. The study was approved by the Ethics Committee of the Otto-von-Guericke University Magdeburg. Parts of the data used here were also used by a recent paper that focused on population receptive field mapping (Liu et al., 2021).

### 2.2 Dataset and experimental design

#### 2.2.1 MRI Scanning

UHF-MRI data were recorded at a whole-body 7 Tesla Siemens MRI scanner in Magdeburg (Siemens Healthcare, Erlangen, Germany) using a 32 channel Nova Medical head coil. First, a whole-brain MP2RAGE sequence with the following parameters was acquired: Voxel resolution: 0.7 mm isotropic, 240 slices, FoV read: 224 mm, TR = 4800 ms, TE = 2.01 ms, TI1/2 = 900/2750 m, GRAPPA 2, sagittal orientation. The additional 7T MP2RAGE sequence with an isotropic resolution of 0.7 mm that was also acquired was used for a different study. Before acquiring the functional scans, shimming was performed and two volumes using echo-planar imaging (EPI) with opposite phase-encoding (PE) polarity were acquired. Then, functional gradient-echo (GE) EPI sequences with the following parameters were acquired: Voxel resolution: 1 mm isotropic, FoV read: 192 mm, TR = 2000 ms, TE = 22 ms, GRAPPA 4, interleaved acquisition, 36 slices. The same sequence was used for all functional tasks (see below). 3T-MRI data were acquired on a separate day at a Philips 3T Achieva dStream MRI scanner (at the in-house facility at the Leibniz Institute for Neurobiology (LIN)), where a standard structural 3D MPRAGE was acquired (resolution: 1.0 mm x 1.0 mm x 1.0 mm, TI = 650 ms, echo spacing = 6.6 ms, TE = 3.93 ms, flip-angle= 10°, bandwidth = 130 Hz/pixel, FOV = 256 mm×240 mm, slab thickness = 192 mm, 128 slices).

#### 2.2.2 Stimuli and Design

Participants were stimulated on their individual digits using a piezoelectric stimulator (Quaerosys, Schotten, Germany). We used five MR-compatible and independently controlled piezoelectric modules for tactile stimulation to the right hand’s D1-D5, where D1 corresponds to the thumb, D2 to the index finger, D3 to the middle finger, D4 to the ring finger, and D5 to the little finger of the right hand of younger and older adults. Stimulation was applied while they were lying in the 7T-MRI scanner. Each finger was attached to one module using a metal-free, custom-build applicator that could be easily adjusted to different hand and finger sizes (i.e., each module could be independently moved within the applicator until the pins in the stimulator were positioned under each fingertip). The stimulator (see Figure 1A) had pins arranged in a 2 × 4 array in the proximo-distal axis of the finger, covering a skin area of 2.5 × 9 mm^2^. The vibrotactile stimulation was applied to the fingertips at a frequency of 16 Hz with only two pins rising at a given time to prevent adaptation (Schweizer et al., 2001). The frequency was a continuous sinusoidal function with the intensity adjusted to 2.5 times the individual tactile detection threshold for each subject and finger. Tactile detection thresholds were acquired on a separate day before scanning. The mean detection threshold used for older adults was 1.37 ± 0.07 g, and for younger adults, it was 0.80 ± 0.04 g (Liu et al., 2021).

**Figure 1:**
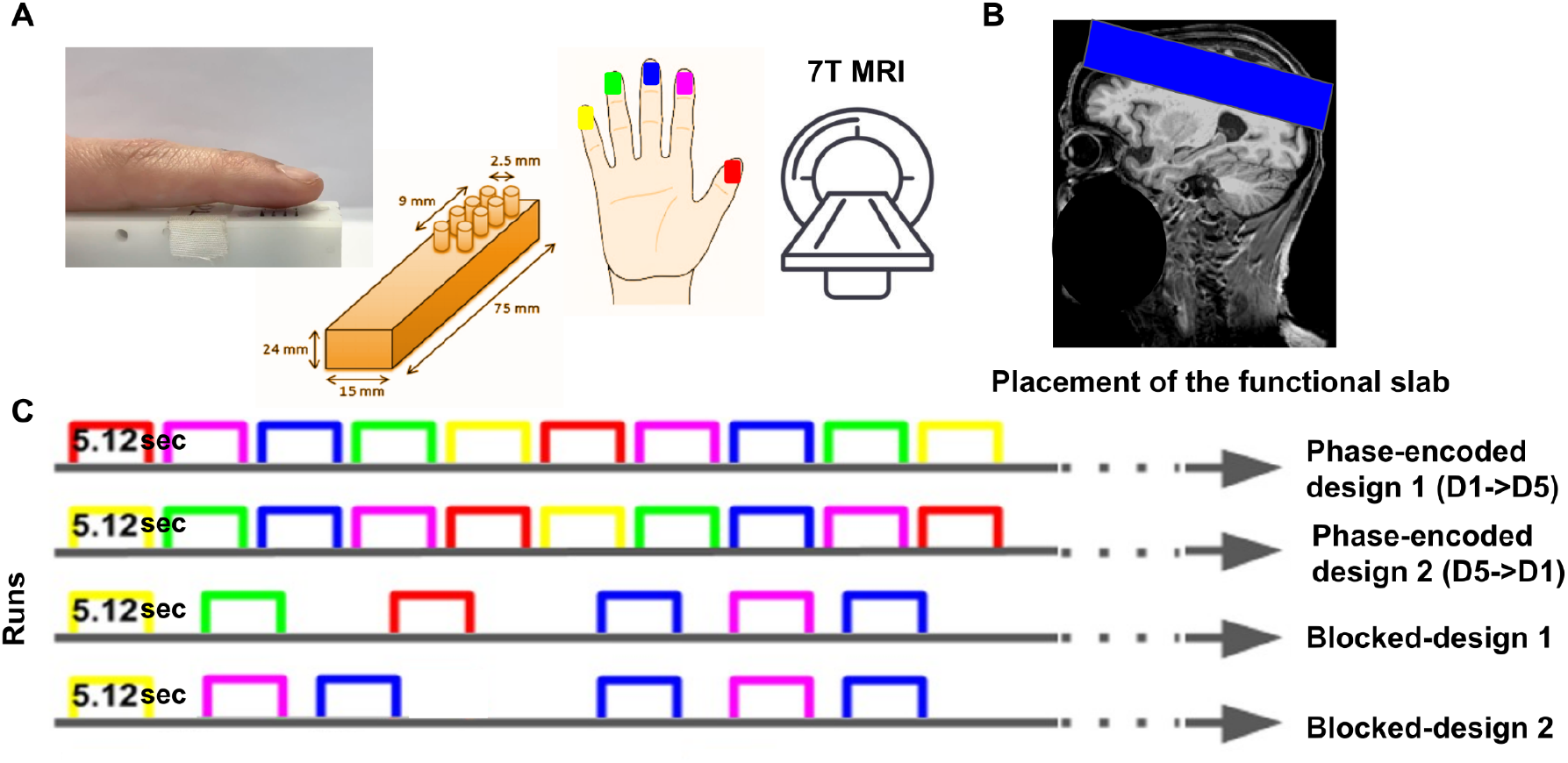
Experimental design. Within the 7T-MRI scanner, a piezoelectric stimulator (see **A**) was used for fingertip stimulation. **B:** The blue area shows the placement of the slab image at an example participant’s T1-weighted image. **C:** Each row represents one run with the first two rows representing phase-encoded designs (Forward: D1 to D5, Reverse: D5 to D1) and the last two rows representing blocked-designs (each row represents one run). Each colour represents one finger of the right hand (colour code in **A**).

There were two phase-encoded runs and, consecutively, two blocked-design runs. The phase-encoded runs consisted of 2 runs of 20 cycles each, with a duration of 25.6 seconds per cycle. On each run, each fingertip received stimulation for 5.12 seconds and was repeated 20 times. There was no delay between the end of one stimulation on one finger and the start of the next at a different finger. For each run, the stimulation was applied either in forward (D1 to D5) or reverse (D5 to D1) order. Forward and reverse orders were randomized across participants and age groups (i.e., half of the participants of each age group, and of each gender started with a forward run). Each run consisted of 256 scans (512 sec for TR = 2 seconds) and lasted for 8 minutes and 31 seconds with the same number of intervals in each run (15 in total), there was 1-minute rest between runs and no gap between repetitions. The order, forward (D1 to D5) or reverse (D5 to D1) order was chosen because the same design has been previously used and it was found to be very sensitive to detect precise topographic finger maps in area 3b (Kuehn et al., 2018).

For the blocked-design, there were 6 conditions: Stimulation to D1, D2, D3, D4, D5, and a rest condition. The stimulation protocol was similar to the phase-encoded design, i.e. each finger was stimulated for 5.12 seconds with the same frequency and intensity as described above. In contrast to the phase-encoded design, the order of finger stimulation was in a pseudo-random order, which was randomized in a way that no finger was stimulated more than twice in a row. There was a 2 seconds rest in 70% of the trials and a 6 seconds rest in the remaining 30% between every two subsequent stimulations. Every finger was stimulated 10 times. Each run comprised 210 scans, which lasted for 6 min and 56 sec. The blocked-design paradigm was repeated twice as well. The phase-encoded runs were used for the analyses using SRM and blocked-design runs were used as a localizer for defining the ROI.

During both phase-encoded and blocked-design runs, there were small gaps of 180 ms inserted into the 5.12 seconds stimulation intervals to motivate participants to focus on the stimulation (Schweisfurth et al., 2014).These gaps were randomly inserted within one stimulation block. It was the task of the participants in all 4 runs to count the number of gaps throughout the block and to verbally report the number after the block was finished. Gaps were pseudo-randomized in a way that each finger received the same number of gaps throughout the experiment. Counting the gaps ensured high attention of the participants towards the stimulus.

### 2.3 Preprocessing

Functional MR data acquired at 7T can be corrupted by motion artifacts and geometric distortions due to field inhomogeneity. To counter distortions, two opposite PE polarity EPIs were acquired prior to the functional scan. To perform distortion correction of both EPIs with opposite PE polarity, a point spread function (PSF) mapping method was applied (In et al., 2016). A weighted combination of the two distortion-corrected images was incorporated to maximize the spatial information content of the final corrected image, because the amount of spatial information differs between the opposite PE datasets. The EPI-images of the functional blocks were motion corrected to time point t_0_ = 0, and the extended PSF method was applied to the acquired and motion-corrected images to perform geometrically accurate image reconstruction. This was followed by slice-timing correction using SPM8 (Statistical Parametric Mapping, Wellcome Department of Imaging Neuroscience, University College London, UK).

### 2.4 Decoding Analyses

The 1st level fixed-effects models were computed separately for every subject using the general linear model (GLM) implemented in SPM8. This analysis was performed on the two blocked-design runs (see **Figure 1C**). We modeled five regressors, one per digit (D1, D2, D3, D4, D5 stimulation). Then, five linear contrast estimates were computed (e.g. contrast [4 −1 −1 −1 −1] for touch to D1). No group alignment was performed; therefore, the anatomical and functional data were neither smoothed nor normalized. It is important to note that splitting the data into two phases (blocked-design and phase-encoded design) allowed us to have independent datasets for defining the ROIs (i.e., the fingers) and training the SRM. Using a blocked-design with randomized stimulation sequences allowed an optimal localization of each individual finger in each participant. The phase encoded design was chosen for the SRM, because the model requires the stimulation order to be the same for each participant, and using one forward- and one backward-run ensured that we controlled for effects of stimulation order. Therefore, for the first analysis scheme (see section 3.2 for more details), the independently acquired blocked-design data was used for masking the top n-number [500, 1000, 1500, 2000, 2500, 3000] of significant voxels from the phase-encoded data.

For the whole-brain region of interest (ROI) based analysis scheme, the focus was on investigating the representations in different BAs, more precisely, to investigate seven BAs covering the sensorimotor system (BA1, BA2, BA3a, BA3b, BA4a, BA4p, BA6) in the hemisphere contralateral to the stimulation (i.e., right-hand stimulation, left hemisphere). Freesurfer (version-v6.0.0) based parcellation was performed using the command “recon-all” on the T1 anatomical maps. Note that both the structural data (3T MPRAGE data) and the functional data (7T EPI) had the same resolution (i.e., 1 mm isotropic) and could therefore be mapped onto each other. In order to co-register the 7T functional and the 3T structural images, a function-to-structure manual rigid co-registration was performed using ITK-snap (version 3.8) and Advanced Normalization Tools (ANTs version 2.1). ITK-snap was used to generate a transformation matrix, which was used to transform the functional data using ANTs. Rigid co-registration was performed because there were no morphological differences between the reference and the source. To maintain homogeneity, the top 500 voxels were selected from each region to perform the analysis across age groups and across different brain regions. This number was selected based on the highest stimulus decoding accuracy obtained for 500 voxels as compared to the other numbers. For columnar-based analyses, we focused on the precise functional representation of the somatosensory finger maps that exist in BA1 and BA3b in the contralateral (left) hemisphere.

#### 2.4.1 Shared Response Modeling (SRM) and digit and age classification in different BAs

##### 2.4.1.1 The Shared Response Model

The underlying concept of SRM is to determine a shared, lower-dimensional representation of the stimulus-response across subjects who were presented with the same stimulus during scanning (here synchronized finger stimulation). Please note that SRM can, however, also be used to detect differences between individuals and groups, which corresponds to the approach used here. Shared Response Modeling and classification were performed in two separate leave-one-subject-out procedures. We separated SRM calculation from classification to reduce the computational costs. Importantly, no data from the left-out subject were used to estimate the SRM or classifier model. More precisely, we applied robust SRM (Turek et al., 2018) to find the shared response of all subjects except the left-out subject using the first of the two runs with phase-encoded design. This resulted in the matrices *W_j_* ∈ ℝ_*N×K*_, *j* = 1…*m* – 1, which we refer to as subject-specific bases, where *N* is the number of voxels, *K* is the number of shared responses and *m* is the number of subjects. These bases can be considered mappings of individual topographies where the BOLD signal *X_j_* ∈ ℝ_*N×t*_, is approximately *X_j_* ≈ *W_j_R* with *R* ∈ ℝ^*K×t*^ (see **Figure 2**). The basis *W_m_* of the left-out subject is then determined using the BOLD signal *X*_*m*,1_ of the first of two runs with phase-encoded design and the shared response *R* as a template. Finally, the BOLD signal *X*_*m*,2_ of the second of the two runs with phase-encoded design was projected to the *K*-dimensional feature matrix *R_m_* ∈ ℝ^*K×t*^ using the basis *W_m_*. The individual matrices *R_m_* served as input data for the classifier in the functionally aligned feature space. This step was repeated for all subjects. The leave-one-subject-out procedure was performed separately for the two age groups. These separately trained shared spaces were used to differentiate the two age groups rather than aligning.

**Figure 2:**
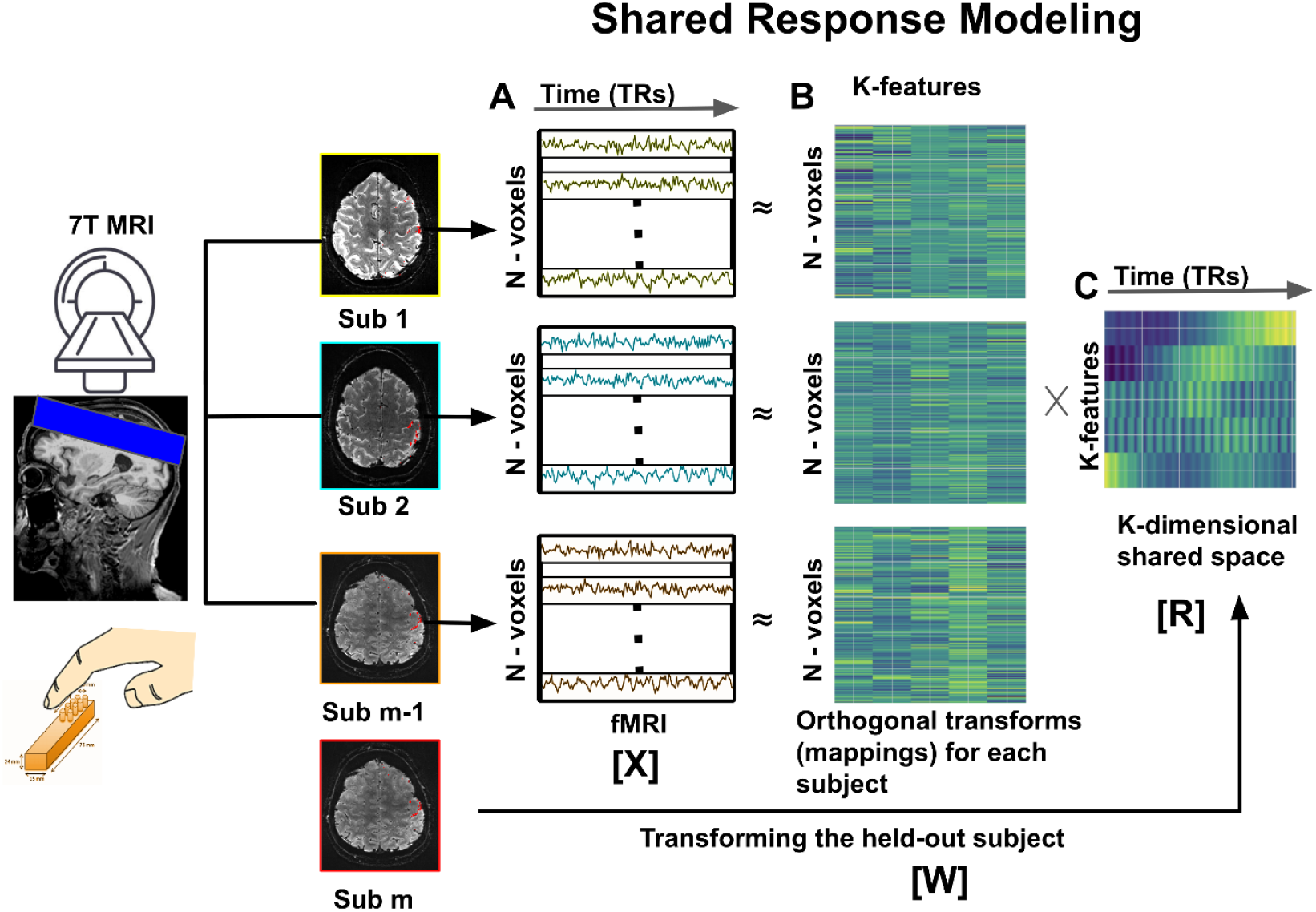
Workflow of Shared Response Modeling (SRM). **A:** The model assumes that all participants receive the same stimulation during the scanning; therefore, phase-encoded designs were used for analyses (see **Figure 1C** for details). The matrix [X] represents the subjectwise BOLD signal. **B**: The matrix [W] with N voxels by *K* features is crucial for participant-specific mapping, and is also referred to as ‘subject-specific basis’ in the text. **C**: The model captures the underlying shared variance with *K*-dimensional shared space representation across subjects for all samples represented by matrix [R].

##### 2.4.1.2 Digit classification

For digit classification, we used each scanned volume of the projected reduced feature space as training sample where the corresponding stimulated digit was set as class label, i.e. the feature space was *K*-dimensional. A linear support vector machine (SVM) classifier was trained and tested in a leave-one-subject-out cross-validation approach. The achieved decoding accuracy we consider as an estimate of the alignment capability of SRM using different selection methods of voxels (for BAs and columns, see **Figure 3A and Figure 5D,E** respectively). Digit classification was carried out separately for the group of younger and older adults (i.e., two leave-on-subject-out cross-validations were performed, see **Figure 3A, Figure 4A**). The chance accuracy (i.e. Ac_chance_) for this decoding analysis is 0.2, given 5 fingers were used for classification.

**Figure 3.**
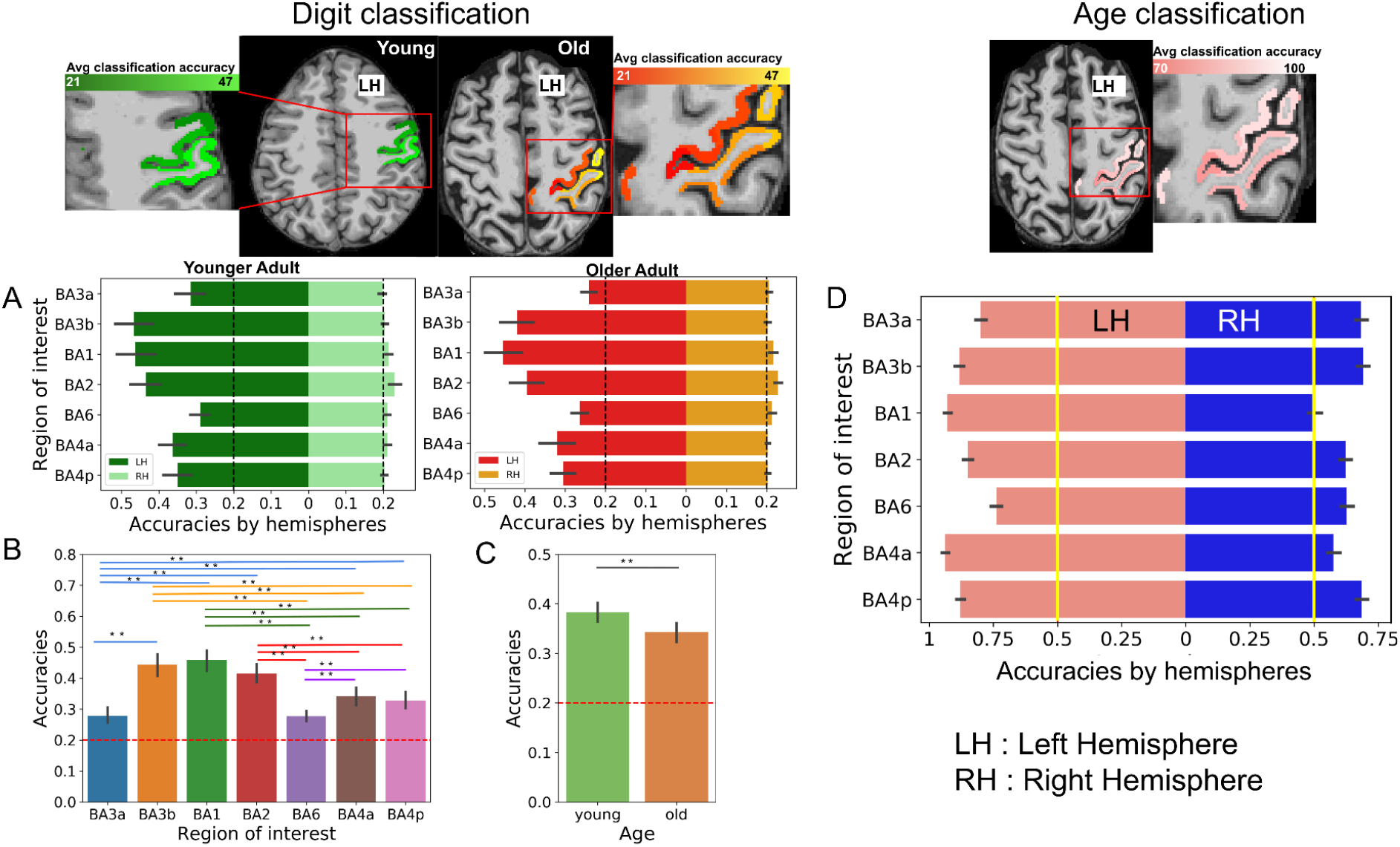
Sensory Finger Decoding in Different BAs in Younger and Older Adults. **Digit classification** (*left*): **A:** Bar plots show ROI-based decoding accuracies for the left and right hemispheres obtained by responses to digit stimulation in younger and older adults. Average accuracies are mapped over the brain at the top row (younger (green) and older (red) adults). The black dotted line indicates chance level for finger classification (5 classes, chance accuracy (i.e. Ac_chance_) = 0.2)). **B:** Significant main effect of ROI in digit-discriminating decoding accuracies. Asterisks represent significant differences at the p<0.05 level, and double asterisks represent significant differences at the p<0.005. Colours indicate the different regions of interest. **C:** Significant main effect of age group in digit decoding accuracies, (younger (green) and older (orange) adults). Double asterisks represent significant differences at the p<0.005 level. **D:** Comparison between age group decoding accuracy between right (ipsilateral) and left (contralateral) hemispheres using SRM. Note that tactile stimulation was provided to the fingers of the right hand. Error bars represent the standard error over the accuracy scores obtained using SVM. The red dotted line indicates chance level for age classification (2 classes, Ac_chance_ = 0.5).

**Figure 4:**
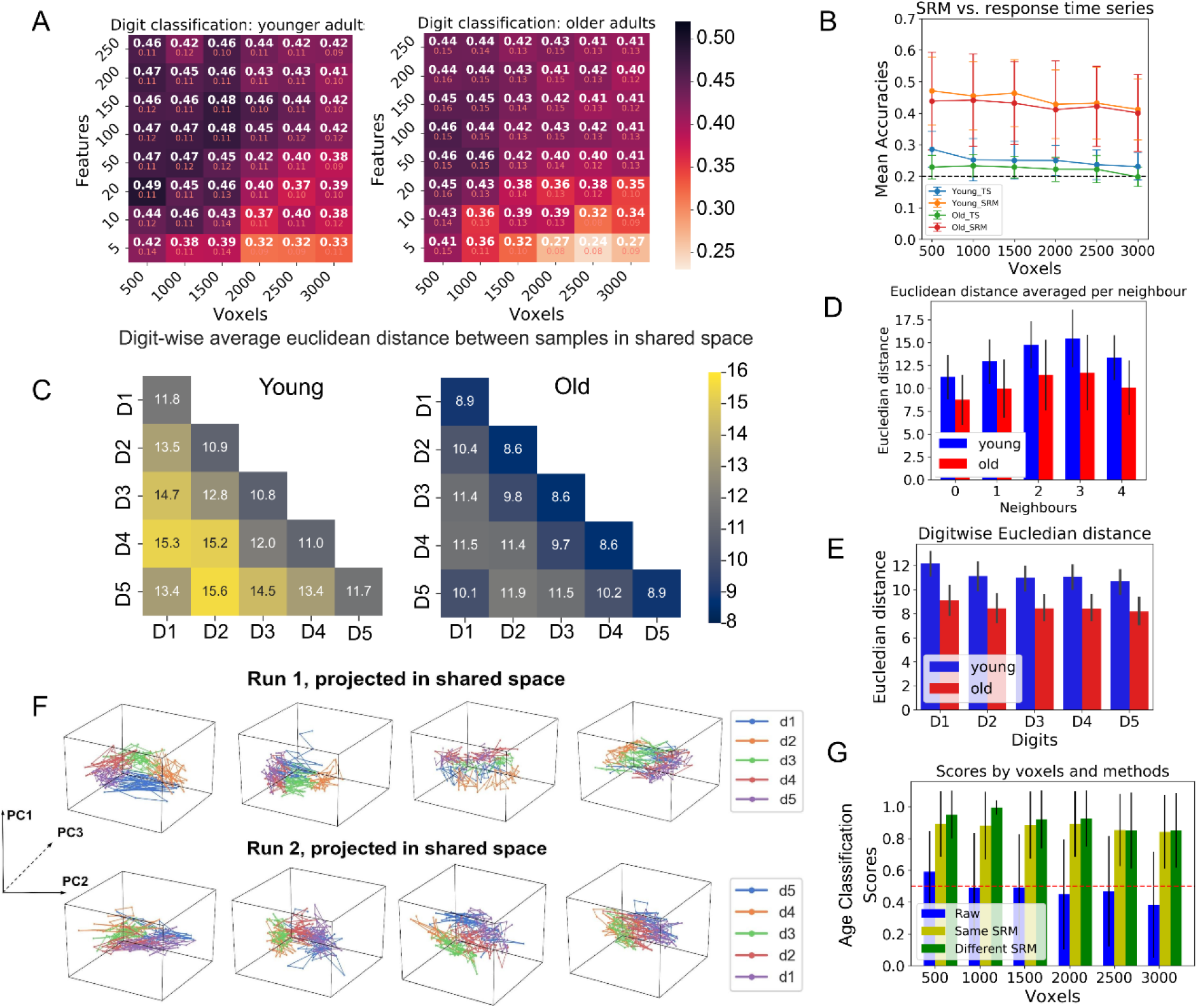
Age-based SRM Classification. **A:** Heatmaps show mean accuracies and standard deviations of different feature numbers and numbers of voxels. Accuracies were obtained by leave-one-subject-out cross-validation using SVM with a linear kernel. **B:** Line plots show the unaggregated time-series-based (*green* for younger and *blue* for older adults) and SRM-based (*orange* for younger and *red* for older adults) digit decoding accuracies (with k = 200); error bars show the standard deviation. **C:** Average Euclidean distances between samples in shared space generated using SRM with K = 5 features for younger and older adults. The low values in the diagonal and the higher values in the off-diagonal show that similar digit responses are closer to each other and decrease gradually along the first, second, third, and fourth neighbour. **D:** Bar graphs show average Euclidean distances as in **C** but ordered along the dimension of neighbour (first, second, third, fourth neighbour, note that there are more first then second neighbours, more second than third neighbours, and more third than fourth neighbours) and with a direct comparison between younger (*blue*) and older (*red*) adults. Error bars indicate standard deviations. **E:** Bar graphs show average Euclidean distances for younger (*blue*) and older (*red*) adults digit-wise. Error bars indicate standard errors of the mean. **F:** Top three principal components obtained by PCA of run “D1 to D5” and run “D5 to D1” projected into shared space. The three axes represent the principal components of the shared response in the 3D representationsl space. Digits are arranged circularly in 3D space for 4 different example subjects. NOTE: PCA was only performed for visualization purposes this is not part of any of the analyses. **G:** Bar graphs show mean age decoding accuracies of the fMRI time series unaggregated (*blue*), the accuracy of the projected shared space data when both age groups were trained using the same shared response model projected in a cross-validated way (*yellow*), and when different shared spaces were learned for younger and older adults, respectively (*green*). Accuracy scores were obtained by using linear SVM.

##### 2.4.1.3 Age classification

For age classification, we used each scanned volume after projection to the reduced feature space as training sample, i.e. the feature space was *K*-dimensional, but this time, the age group was set as class label, irrespective of the simulated finger. In contrast to digit classification, we involved the previously calculated shared response features of all subjects, i.e. of both age groups, for age classification. To achieve balanced training sets and to provide a reliable estimate of accuracy, we left out two subjects, one of each group, in the cross-validation procedure and repeated this step for each possible combination of selecting one subject from the younger and one subject from the older group. This resulted in a distribution of 171 accuracies from which we determined the mean and standard deviation. Note that the number of samples per class is greater compared to digit classification and the amount of training data is greater. The chance accuracy Acchance for this decoding analysis is 0.5, given two age groups were used for classification (see **Figure 4G, Figure 3D**).

#### 2.4.2 Columnar-Shared Response Modeling (C-SRM)

Sensory and motor cortices can be divided into columnar units, which helps to detect fine-grained topographic maps with high precision (Huber et al., 2020a; Yacoub et al., 2008; Yang et al., 2019). We here aim to identify the number of columns required to effectively decode the SI generated responses, i.e., the topographic map of the fingers in younger and older adults. Note that because each finger was stimulated by 16 different combinations of pins (see **Figure 1A**), the smallest computational unit may not be the representation of one finger but the representation of one or more pins touching the skin. We focus again on the hemisphere contralateral to the stimulation (i.e., right hand stimulation, left hemisphere), and on the areas that offer a fine-grained representation of touch (i.e., BA1 and BA3b). First, we divide the two ROIs BA1 and BA3b into different numbers of columns (ranging from 10 to 400 approximately equi-volume columnar units, where the maximum number of possible columns would be represented by the number of voxels) using the software package LAYNII2 (i.e. Version 2.0 of Laynii) (command: LN2_COLUMNS) (Huber et al., 2021). This step groups the thresholded voxels into different columnar units. This command from the package implements a method (see, Huber et al., 2021) that generates approximately equal volume columns in the T1-MRI image by using the middle gray matter segment as the input, which is generated by LN2_LAYERS (i.e. the cortical columns were defined based on gray matter anatomy after performing structure to functional registration). Then, we took the mean of all the thresholded voxels within a columnar structure to compute a matrix: columns by time-series (see **Figure 5C**). This was done for both phase-encoded runs and across all three ROIs for all subjects. Then, this matrix was used to train the SRM in a cross-validation way, as described in the previous section. Linear SVM-based leave-one-subject-out decoding analyses were then performed separately for the different number of columns to verify that the model can successfully capture the digit stimulus-based features, and to determine the number of columnar divisions that best captures these features. In this way, we modeled the functional activation patterns to find the optimal number of columns, which were initially generated based on anatomy of the cortex.

**Figure 5.**
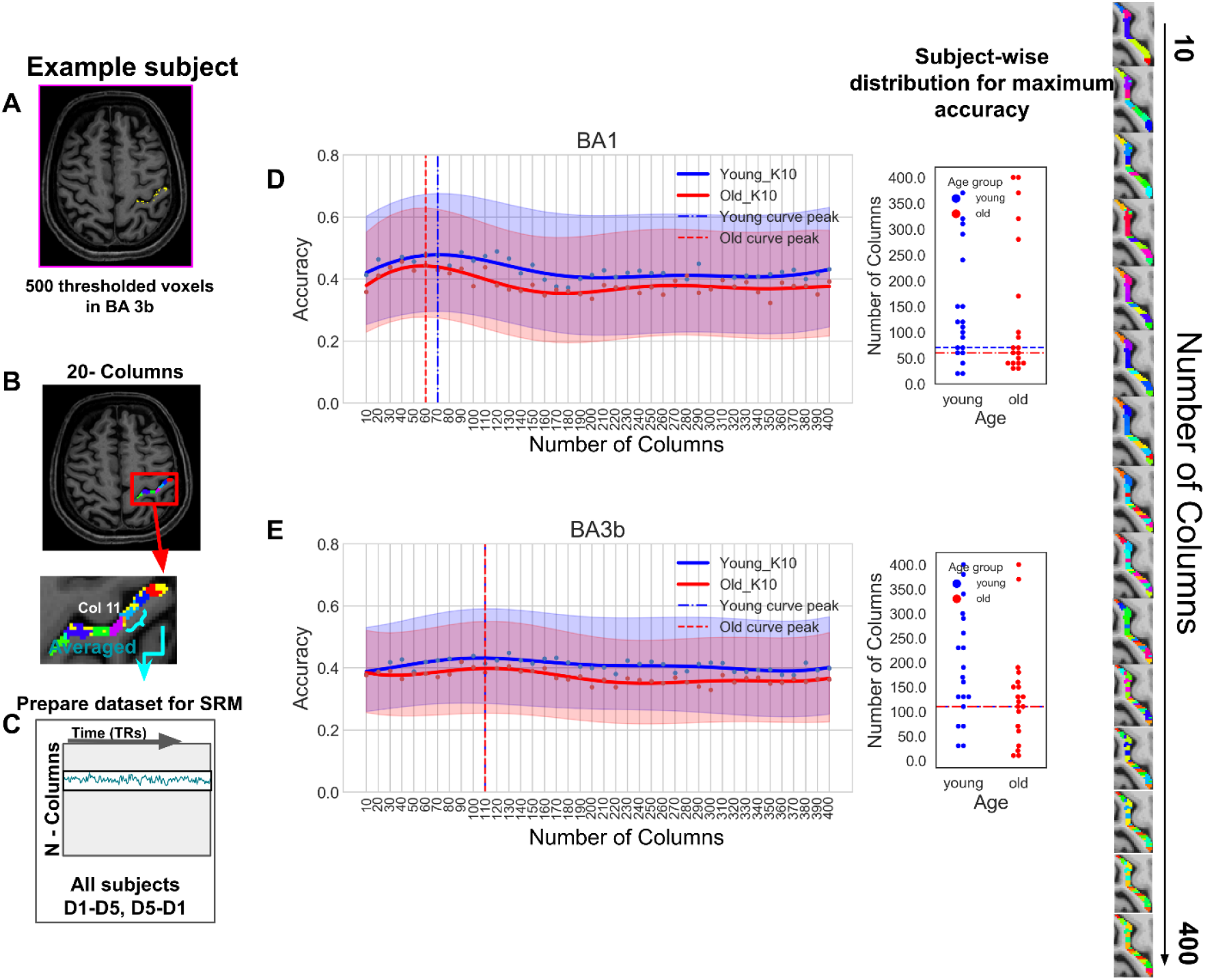
Column-Based SRM (C-SRM) **A, B:** An example subject’s anatomical data showing different numbers of columns in BA3b, and a schematic overview of how the shared response space is generated for N different numbers of columns. **C:** Dataset matrix represents averaged time-response of voxels present in each column for both run types (D1 to D5, D5 to D1). **D-E:** Means (*solid lines*) and standard deviations (*semi-transparent spread*) of digit classification scores over a range of columns and different age groups; 10 features were used to train the SRM; the plot is a smoothed curve obtained by Gaussian smoothing for a window-size of 6. **D:** Mean and standard deviation of digit decoding scores in ROI BA1 shown for different numbers of columns; the different colours indicate the age groups (younger participants: *blue*, older participants: *red); vertical dashed lines* indicate curve maxima); a subject-wise distribution with the number of columns at which the maximum accuracy was achieved is shown in the adjacent plots (*right*). **E:** same as (**D**) but for the ROI BA3b.

### 2.5 Statistical Analyses

A two-way ANOVA with the factors age (younger, older) and ROI was calculated on the digit classification accuracies to test the influence of both factors, and the interaction between both factors, on the digit classification scores in the ROI-based analyses. For significant main effects or interactions, post hoc independent sample t-tests were computed. The statistical tests were performed in Python using the ‘statsmodels’ package. An alpha level of p<0.05 was used to test for the significant main effects and interactions, and for significant post hoc test results. We also performed a permutation test to evaluate the decoding model. For this, we permuted the target (i.e. age labels) to generate randomized data and computed the ‘p’ (p-value) against the null hypothesis that the shared-space features and age labels are independent.

## 3 Results

### 3.1 Finger-specific sensory decoding in different BAs

Different brain areas respond differently to somatosensory stimulation, in particular with respect to signal specificity. In this section, we use SRM to describe the hierarchical encoding of somatosensory stimuli in higher- and lower-order somatosensory and motor cortices of the hemisphere contralateral to the stimulation (right hand stimulation, left hemisphere). To this aim, we compared sensory decoding accuracies across different Brodmann areas (BA1, BA2, BA3a, BA3b, BA4a, BA4p, and BA6). In both age groups, BA1 and BA3b revealed the highest mean decoding accuracies (BA1: accuracy younger: 0.46 ± 0.12, accuracy older: 0.45 ± 0.10, BA3b: accuracy younger: 0.47 ±0.12, accuracy older: 0.42 ± 0.10), followed by BA2 (accuracy younger: 0.43 ± 0.09, accuracy older: 0.39 ± 0.10), BA4a (accuracy younger: 0.36 ± 0.08, accuracy older: 0.32 ± 0.10) and BA4p (accuracy younger: 0.35 ± 0.09, accuracy older: 0.31 ± 0.07). Lowest accuracies were revealed for BA6 (accuracy younger: 0.29 ± 0.06, accuracy older: 0.26 ± 0.06) and BA3a (accuracy younger: 0.32 ± 0.09, accuracy older: 0.24 ± 0.04). Given that BA3b and BA1 are known to contain the most precise topographic finger maps in younger adults (Pfannmöller et al., 2016), our methodological approach supports this finding and additionally reports this for the first time for older adults.

We were additionally interested whether within this given processing hierarchy, age-differences would be apparent. A two-way ANOVA with the factors age (younger, older) and BA (BA 1, 2, 3a, 3b, 4a, 4p and 6) revealed a main effect of age (F(1) = 14.13, p = 2.12e-04) and a main effect of BA (F(6) = 27.78, p = 2.17e-25) over digit classification accuracies, but no significant interaction effect between BA and age (F(6) = 0.49, p = 0.815). The main effect of age was due to older adults showing generally worse decoding accuracy compared to younger adults across different BAs (average accuracy, younger: 0.38 ∓ 0.11; older: 0.34 ∓ 0.10).

The main effect of BA was due to (i) significantly lower accuracy of BA3a compared to BA3b (BA3a: 0.28 ± 0.08, BA3b: 0.44 ± 0.11, t(74) = 7.52, p = 1.02e-10), BA1 (BA1: 0.46 ± 0.11, t(74) = 8.09, p = 8.61e-12), BA2 (BA2: 0.41 ± 0.09, t(74) = 6.89, p = 1.59e-9), BA4a (BA4a: 0.34 ± 0.09, t(74) = 3.23, p = 0.002), and BA4p (BA4p: 0.33 ± 0.08, t(74) = 2.66, p = 9.49e-03); (ii) significantly higher accuracy of BA3b compared to BA4a (t(74) = 4.43, p = 3.16e-05), BA4p (t(74) = 5.18, p = 1.87e-06), and BA6 (BA6: 0.28 ± 0.05, t(74) = 8.44, p = 1.93e-12); (iii) significantly higher accuracy of BA2 compared to BA4a (t(74) = 3.51, p = 7.68e-04), BA6 (t(74) = 7.94, p = 1.66e-11), BA4p (t(74) = 4.31, p = 5.02e-05).

Taken together, these data show that, whereas the processing hierarchy is preserved across age groups, older adults show generally worse decoding accuracies across different BAs.

### 3.2 Fine-grained SI topography in different age groups

Next, we used SRM to compute across-run digit and age classification to more specifically target the question of whether SRM can be used to describe the above reported age-related changes in the topographic SI architecture in more detail. Note that in the analyses reported above, we show generally lower decoding accuracies as a main effect of age across all BAs, whereas here, we explore the *features* that drive these differences in more detail. We first tested the digit decoding capability of the model. By doing so, we investigated if the number of statistically significant voxels included in the analyses has any influence on the overall decoding accuracy of the digits. This was tested for younger and older adults separately. We found that a higher number of voxels resulted in a higher dimensional feature space capturing the shared response and explaining the shared variance. For example, in the younger age group, to obtain an average accuracy of 0.42 ± 0.14, 500 voxels needed 5 number of features in the shared space whereas 3000 voxels needed 100 number of features in the shared space to obtain the similar decoding accuracy (see **Figure 4A**). This was similar for younger and older adults (in spite of the above mentioned lower decoding accuracies in older compared to younger adults). For further analyses, the optimal number of voxels for the ROI-based analysis was chosen (i.e., n_Voxels_ = 500), with the lower number of features (k =10) explaining most of the shared variance across participants.

Using these parameters, we then tested the performance of the SRM against the unaggregated functional time series based tactile stimulus decoding, and age-group decoding using leave-one-subject-out cross-validation and SVM classification. Here, we first compared the tactile stimulus decoding accuracy in younger and older adults when separately modeled in a shared space using time-series-based classification. We observed that the SRM-based stimulus decoding outperforms the unaggregated time-series-based decoding (e.g. average accuracy scores at 500 voxels selected for younger SRM (k = 200): 0.47 ± 0.11; younger unaggregated time-series: 0.29 ± 0.06, older SRM: 0.44 ± 0.16; older unaggregated time-series: 0.23 ± 0.04), hence confirming that SRM can successfully capture the shared vibrotactile information across subject groups (see **Figure 4B**). Even in the case of age-based decoding analysis, SRM-based decoding performs better than the unaggregated time-series-based decoding (see **Figure 4G,** blue bar) with scores around the chance accuracy of 0.5, whereas the average accuracy achieved using SRM is 0.87 ± 0.02 and 0.91 ± 0.05 (see **Figure 4G**). The bar plot (yellow, see **Figure 4G**) represents the average age decoding accuracy of 0.87 ± 0.02 when a single shared space was used to capture the variance across subjects; the green bar represents the average age decoding accuracy of 0.91 ± 0.05 when separate models were trained in order to learn their respective shared feature space. In both cases, the accuracies obtained were above chance level (i.e. above 0.5). However, when separate models were used to learn the shared feature space, the mean decoding accuracy was higher than when a single model was used. This is because group-specific differences exist, and separate models learn the features shared within a group.

We also performed a permutation test to evaluate the significance of the accuracies obtained by two-fold cross-validation. The dependency between the unaggregated time-series based feature and age labels was low (with p = 0.068) whereas the relationship between the features and age-group labels when the same shared space was used for both the age-groups was high (with p = 0.001). The same was true when separate shared spaces were used to capture the group-specific feature (p = 0.001). This indicates that the age classification accuracies were not obtained by chance.

Given this approach, we then computed digit classification and Euclidean distances to define age-specific changes that may target only specific fingers, or only specific neighbours (see **Figure 4C**). Euclidean distance was measured in the response-driven shared feature space with k = 5 as the number of feature dimensions for training the SRM, we here assumed that each dimension represents a digit. This is relevant, because prior research showed that older adults show reduced spatial distances specifically between area BA3b’s finger representations of D2 and D3, and show relatively less sensory confusion between third-neighbour fingers compared to younger adults (Liu et al. 2021). For both younger and older adults, the diagonal values are smaller than the off-diagonal values, which suggests, as expected, that the sample points corresponding to the same digit are more similar than the neighbouring digits in the shared feature space. In addition, an overall difference in the similarity measure using the Euclidean distance can be observed across age groups with the average Euclidean distance being higher in younger adults as compared to older adults (see **Figure 4C, D**). This indicates that SRM captures the stimulus-specific brain activity that is shared across the participants rather than being affected by the spatial mismatch of functional topographies.

Taken together, the above described results show that i) digit decoding is lower for older compared to younger adults, ii) the sample points corresponding to different digits are arranged more closely to each other in the shared space for both younger and older adults (see **Figure 4D**), and iii) the Euclidean distance measure for older adults is lower compared to younger adults across different digits and neighbours.

### 3.3 Columnar-Shared Response Modeling (C-SRM)

Finally, we used the above outlined methodology to investigate columnar units in SI. As outlined in the introduction, sensory and motor cortices can be divided into columnar units, which helps to detect fine-grained topographic maps with high precision (Yacoub et al., 2008; Yang et al., 2019; Huber et al., 2020a). We here aim to identify the number of columns required to effectively decode the SI generated responses, i.e., the topographic map of the fingers in younger and older adults. Because 16 pin combinations were used to stimulate each finger, the data could potentially identify columnar units that are smaller than one finger representation. We divided the cortex into different numbers of columns, and plotted the results against the respective decoding accuracies, smoothed those data and detected the individual peak to identify the columnar size with best accuracy.

Across different columnar divisions, mean accuracies were higher in BA1 compared to BA3b in both younger and older adults (see **Figure 5D,E**). Interestingly, we observe that the average column size that best predicts the digits were different for younger and older adults in BA1 but not in BA3b. Please note that columnar size here does not relate to anatomical cortical columns, but represents a statistical unit that was derived based on our functional modelling results. The question how these units relate to structural units in S1 is not answered here. In addition, the different columnar sizes in younger and older adults could be driven by both higher precision and reduced noise. In BA1, the finger map of younger adults was best represented by more columns compared to older adults (BA1: younger: 70 (average columnar size: 44.26 ± 5.06 mm^3^), older: 60 (average columnar size: 45.47 ± 5.55 mm^3^); whereas it was the same columnar number in BA3b (both 110 for younger (average columnar size: 28.94 ± 3.51 mm^3^) and older (average columnar size: 24.49 ± 2.40 mm^3^) adults, note for example that for a columnar size of 27mm^3^, the dimensions are 3×3×3mm).

Taken together, we here combine SRM with columnar analyses (a method that we introduce as C-SRM) to offer a statistical approach to calculate the number of columns that best represents finger differentiation in different BAs of SI. We show that in BA3b, 110 columns best represent the smallest unit of finger differentiation in younger and older adults, whereas the columnar number is higher for younger adults in BA1, indicating smaller computational units in younger adults. It should be noted that the term ‘column’ here represents a computational unit derived from functional analyses rather than a structural architectonic feature of the system.

## 4 Discussion

Here, we used shared response modeling (SRM) and introduced columnar-SRM (C-SRM) to describe architectural features of somatosensory finger representations in younger and older adults. We show that the somatosensory processing hierarchy is not significantly different between younger and older adults, as denoted by highest digit-classification accuracies in BA1 and BA3b, followed by BA2, BA4a, BA4p, BA6, and finally BA3a. We also show that digit-classification accuracy was lower for older adults as compared to younger adults across these different BAs. Please note that these group differences are not driven by differences in tactile sensitivity as the stimulation intensity was adjusted to the individual tactile detection threshold in each individual. We further show that the average Euclidean distance across sample points in the feature space is lower for older adults as compared to younger adults across digits and finger neighbours. We also introduce a new analysis approach, C-SRM, that we use to detect the optimal columnar size required to efficiently decode finger representations in SI in different age groups. The results show that for finger decoding, a lower number of columnar units (hence a larger columnar size) is optimal in older adults as compared to younger adults in BA1, which indicates larger computational units in older adults’ sensory SI processing. Together, our results allow a better understanding of basic functional features of somatosensory processing and their change with increasing age. The methods used and introduced here also allow modeling shared and distinct features of sensory maps using voxels or columns as input units that allows studying fine-grained functional units and their group-dependent differences in greater detail.

We performed a region-of-interest (ROI) based analysis to determine the hierarchical representations of digits across different somatosensory and motor areas. In previous studies on hierarchical processing of somatosensory information, it has been reported that a Iarge portion of BA2 neurons receive tactile inputs from areas BA3b and BA1, and that tactile neurons in BA2 therefore have more complex and larger population receptive fields and response properties than those in areas BA3b and BA1 (Gardner, 1988). Hierarchical processing is here defined as the level of precision with which touch to a specific finger causes a distinct representation in the cortex. Lower-hierarchical processing therefore assumes higher decoding accuracies (i.e., more precise representation of each finger), whereas higher-hierarchical processing assumes lower decoding accuracies (i.e., less precise representation of each finger, and potentially greater influence of high-level processing). This previous finding was confirmed by our analyses, where both younger and older adults showed highest finger decoding accuracies in BA3b and BA1, followed by BA2.

However, prior research has rarely investigated somatosensory finger representations in motor areas and BA3a, and has also not yet described potential age-dependent differences in the processing hierarchy. Our data reveal that finger-specific representation of tactile input is better (as revealed by decoding accuracy) in motor cortical areas BA4a, BA4p, and BA6 compared to somatosensory BA3a both in younger and older adults. Our data also indicate that there is more finger-specific information represented in anterior compared to posterior primary motor cortex, and compared to premotor cortex both in younger and older adults. In addition, our analyses reveal that whereas decoding accuracy is generally lower in older compared to younger adults, there is no interaction between age and BA. Therefore, age effects were homogeneous across the different areas that process somatosensory information. Our data therefore indicate that even though older adults represent each finger less distinct, the precision hierarchy (i.e., BA1 & BA3b > BA2 > BA4a > BA4p > BA6 > BA3a) is preserved across age groups and there is no area where somatosensory deterioration clearly precedes in healthy aging. Another possibility is that these accuracies can also indicate less noise and the presence of information (Gardumi et al., 2016).

We also used SRM to detect digit- and age-specific differences in SI and compared it with an unaggregated time-series based decoding method. We found that the SRM-based method performs better than the unaggregated time-series based method (quantified on the basis of classification accuracy), and that SRM successfully captures the digit-based and age group-based variances. We also show that when separate models for younger and older adults are used to learn the shared feature space, the mean age group decoding accuracy is higher than when a single model is used for both younger and older adults. The reasons for this difference may be age-based sensory processing differences, e.g. the decline in touch sensitivity at older age, and that a separate model may learn this group-specific difference more accurately.

We also investigated the shared feature space in more detail using the Euclidean distance measure and found that the samples representing the individual digits were arranged close to each other. We performed principal component analysis (PCA) over the projected shared space and observed that the sample points corresponding to the specific digits in a run were clustered together in the 3D component space, and almost formed a ring-like structure reflecting the nature of digit stimulation (note that stimulation order was “D1 to D5” or “D5 to D1”). This is a relevant observation as it indicates that SRM captures the stimulus-specific brain activity that is shared across the participants rather than being affected by the spatial mismatch of functional topographies.

Finally, we here introduce a novel approach for choosing the optimal number of equi-volume columns in which the cortex can be effectively divided in order to describe topographic maps. Note that this method computes decoding analyses on mean voxel activations that were ordered along columnar units before performing the analyses. One precondition for the model was therefore the assumption that meaningful information is present within one columnar unit and across cortical depths (Huber et al., 2017). We named the method C-SRM to highlight the combination between columnar modeling and shared response modeling. C-SRM may be a relevant new method given prior studies used different columnar size to describe topographic maps without providing a reason or statistical approach to justify these numbers. For BA3b, we show that the same number of cortical columns provides highest decoding accuracies for both younger and older adults, even though the analyses were performed separately for each group. This result of 90 equi-volume columns that best represent the number of units that describe topographic finger maps in BA3b is therefore robust and was replicated across age groups. Given this analysis was done in the a priori defined finger map area, this corresponds on average to 22 columns per digit, which is close to the 16 pin combination that were used for stimulating each digit. One may take these data to argue that the SI representation of one pin that was used for finger stimulation approximately gave rise to a distinct cortical signal that was useful for finger-specific decoding. If confirmed by future research, this would be remarkable because it would indicate that columnar mapping can be used to detect the computationally smallest unit in BA3b, which is one population receptive field representing the location of one pin on the skin, with available fMRI resolution.

Interestingly, we also found that the maximum decoding accuracy for older adults was achieved at a lower number of columns for BA1 compared to younger adults. Whereas for younger adults, 70 columns in BA1 best distinguished fingers, for older adults, this was the case for 60 columns in BA1. If we consider the above outlined argument that one column may represent approximately one population receptive field, this aligns with the recent finding that the somatosensory receptive field size for older adults are larger as compared to the younger adults (Liu et al., 2021). One biological mechanism that one could therefore hypothesize to underlie these results is that the increased population receptive field size in older compared to younger humans leads to an enlargement of the columnar units that characterize the system. In prior research, reduced intracortical inhibition has been hypothesized to be one mechanism underlying larger sensorimotor representations in older age (Pleger et al., 2016; Ruitenberg et al., 2019).

Also the visual cortex contains a hierarchy of visual areas. The earliest cortical area (V1) contains neurons which respond to colour, form and motion, and the second visual area (V2) contains a stripe-based anatomical organization, as has been shown in non-human primates (Lund et al., 2003). Neurons in these stripes have been proposed to serve distinct functional roles, e.g. the processing of colour, form and motion. These stripes represent an intermediate stage in the visual hierarchy and serve a key role in the increasing functional specialization of visual areas. In the visual cortex, there are also distinct microstructural features that characterize cortical columns (Erwin et al., 1995). One study investigated ocular dominance columns (ODC) (Chaimow et al., 2017), which is a structure that can be spotted in ex vivo cytochrome oxidase stainings, and that has a distinct functional and connectivity architecture. Neighboring areas in the visual cortex encode different features (in this case different orientations), resulting in a distinct structure. In our case, however, neighboring areas in the finger encode neighboring skin locations, they are therefore most likely structurally less distinct from one another compared to ODCs. Whether our described columnar model also has a structural component in SI and is, for example, accompanied by alterations in neuronal or myelin architectures, remains to be investigated (Doehler et al., 2022).

Taken together, different SRM-based methods were used here to investigate group differences in ultra-high-resolution data, and were shown to be promising as they confirmed previous findings as well as added novel information on the architecture of aging topographic finger maps. In combination with columnar mapping, our study introduces a novel approach to map fine-grained features of sensory space that may otherwise escape detection, such as responses to single pin stimulation. However, it is worth noting that this approach can only be used for synchronized stimulus applications across participants, which requires experimental designs where the stimulation order is the same in all participants, hence not randomized. In addition, highly homogenous stimulation is a requirement, which was possible here by applying highly controlled sensory stimulations to individual fingers. This was ensured in earlier studies by, for example, asking participants to view the same movie (Guntupalli et al., 2016; Häusler and Hanke, 2021). However, when this decoding analysis is applied to other systems or paradigms, such as a motor task using an effective movement tracking system, or in dynamic non-synchronized cognitive tasks, it has to be clarified how subtle inter-subject variances in movement speed or cognitive computation can be accounted for by this method.

## 5 Conclusion

Here, we aggregate UHF-fMRI group data using SRM, and report the model to be successful in capturing digit-specific tactile information in the sensorimotor cortex. We also introduced a novel method, C-SRM, for identifying the optimal number of equi-volume columnar units to achieve the highest somatosensory decoding accuracy that may be a promising approach to combine structural and functional high-precision analyses in the future. SRM also successfully identified age-specific shared variances across age groups (younger and older adults), which makes it a potentially suitable tool for intersubject alignment. The analyses presented here may in the future be used in UHF-fMRI studies to capture the variability across groups and brain regions, and to detect specific responses to small processing units. C-SRM can also be applied to other sensory regions of the brain and reveal information about the optimal number of columnar units.

## 6 Credit authorship contribution statement

**Avinash Kalyani:** Conceptualization, Investigation, Formal analysis, Methodology, Writing - original draft. **Oliver Contier:** Conceptualization, Methodology, Writing - review, and editing. **Lisa Klemm:** Methodology, Writing - review, and editing. **Stefanie Schreiber:** Writing - review, and editing. **Oliver Speck:** Conceptualization, Methodology, Writing - review, and editing. **Elena Azañon:** Methodology, Writing - review, and editing. **Christoph Reichert:** Supervision, Conceptualization, Methodology, Writing - review, and editing. **Esther Kuehn:** Supervision, Investigation, Conceptualization, Methodology, Writing - review, and editing.

## Acknowledgments

This work was funded by the Deutsche Forschungsgemeinschaft (DFG, German Research Foundation): Project-ID 425899996, SFB 1436 and KU 3711/2-1, Project-ID: 423633679. Avinash Kalyani was funded by the Center for Behavioral Brain Sciences Magdeburg - CBBS gefördert durch EFRE, Förderkennzeichen: ZS/2016/04/78113.

## Notes

### Competing Interest Statement

The authors have declared no competing interest.

